# Engineering 6-phosphogluconate dehydrogenase to improve heat tolerance in maize seed development

**DOI:** 10.1101/2020.05.21.108985

**Authors:** Camila Ribeiro, Tracie A. Hennen-Bierwagen, Alan M. Myers, Kenneth Cline, A. Mark Settles

## Abstract

Endosperm starch synthesis is a primary determinant of grain yield and is sensitive to high temperature stress. The maize chloroplast-localized 6-phosphogluconate dehydrogenase (6PGDH), PGD3, is critical for endosperm starch accumulation. Maize also has two cytosolic isozymes, PGD1 and PGD2 that are not required for kernel development. We found that cytosolic PGD1 and PGD2 isozymes have heat stable activity, while amyloplast-localized PGD3 activity is labile under heat stress conditions. We targeted heat-stable 6PGDH to endosperm amyloplasts by fusing the *Waxy1* chloroplast targeting peptide coding sequence to the *Pgd1* and *Pgd2* open reading frames. These WPGD1 and WPGD2 fusion proteins import into isolated chloroplasts demonstrating a functional targeting sequence. Transgenic maize plants expressing WPGD1 and WPGD2 with an endosperm specific promoter increased 6PGDH activity with enhanced heat stability *in vitro*. WPGD1 and WPGD2 transgenes complement the *pgd3* defective kernel phenotype indicating the fusion proteins are targeted to the amyloplast. In the field, the WPGD1 and WPGD2 transgenes can mitigate grain yield losses in high nighttime temperature conditions by increasing kernel number. These results provide insight on subcellular distribution of metabolic activities in the endosperm and suggest the amyloplast pentose phosphate pathway is a heat-sensitive step in maize kernel metabolism that contributes to yield loss during heat stress.

**Significance Statement:** Heat stress reduces yield in maize by affecting the number of kernels that develop and the accumulation of seed storage molecules during grain fill. Climate change is expected to increase frequency and duration of high temperature stress, which will lower grain yields. Here we show that one enzyme in central carbon metabolism is sensitive to high temperatures. By providing a heat-resistant form of the enzyme in the correct subcellular compartment, a larger number of kernels develop per plant during heat stress in the field. This genetic improvement could be included as part of integrated approaches to mitigate yield losses due to climate change.

## Introduction

At current consumption levels, the estimated human population of ~9.7 billion in the year 2050 will require a 50% increase in cereal grain production (1, 2). Maize (*Zea mays L.*) provides 42% of worldwide tonnage and is the most prominent cereal grain crop (3). Cereal yield is expected to diminish due to increased abiotic and biotic stress arising from anthropogenic climate change with heat stress already impacting maize yields in Europe (4). Maize breeding has enhanced yield, grain quality, and abiotic stress tolerance to expand the production range, but tolerance to abiotic stresses is challenging for breeders to improve (5). Higher temperatures impact kernel number and accelerate maize kernel development resulting in lower number and reduced weight of mature kernels (6, 7). Discovering biochemical mechanisms that underlie these physiological responses will enable breeding and genetic engineering approaches to maintain cereal grain yield as global temperatures increase.

The plastidial form of the pentose phosphate pathway (PPP) enzyme 6-phosphogluconate dehydrogenase (6PGDH) is required for normal maize endosperm starch biosynthesis (5) and thus is a candidate to affect yield variation in response to heat stress. Starch is the most abundant storage molecule in maize kernels with increased content directly related to higher yield in hybrids (8). Most of the glucose routed from central metabolism into endosperm starch is cycled through glycolysis and the PPP (9). 6PGDH catalyzes the last step of the PPP oxidative phase (oxPPP), generating ribulose-5-P and NADPH (10). The oxPPP enzymes, including 6PGDH, are found in both the cytosol and plastid stroma. The *pgd1* and *pgd2* loci encode cytosolic 6PGDH (11), and double null mutants do not show any kernel phenotypes indicating that cytosolic 6PGDH is dispensable for kernel development (12, 13). By contrast, null mutations of *pgd3*, which encodes the plastidial form, cause reduced starch accumulation and disrupt embryo development, resulting in severe defective kernel phenotypes (14, 15). A small fraction of *pgd3* mutants can germinate in tissue culture and complete a full life cycle, indicating PGD3 is primarily essential for kernel development (15).

This study demonstrated that plastidial 6PGDH is a factor contributing to yield loss in high temperature stress. Temperature-stable forms of 6PGDH expressed in maize endosperm plastids increased enzyme activity and mitigated the reduction in grain yield that occurred in control plants exposed to elevated temperatures at night.

## Results

### Plastid-localized 6PGDH is solely required for kernel development

Prior phenotypic comparisons of *pgd1-null; pgd2-125* double mutants and *pgd3* mutants were in different genetic backgrounds (14). This experiment was repeated in a consistent genetic background to rule out the possibility that the previous *pgd1; pgd2* line requires less total 6PGDH activity for kernel development and that the double mutant might have a kernel phenotype in other inbred lines. To enable marker assisted selection during backcrossing, the genomic sequences of the *pgd1-null* and *pgd2-125* alleles were determined. The *pgd1-null* allele has a complex insertion-deletion polymorphism in the 3’ end of the open reading frame (ORF). The polymorphism has a 5 bp deletion with a 12 bp insertion and a second 3 bp deletion with an 18 bp insertion. The last 13 codons of the ORF are replaced to code for 37 novel amino acids at the PGD1^null^ C-terminus (**Fig S1A**). The *pgd2-125* allele has a missense mutation of R460T, which also impacts the C-terminal domain (**Fig. S1A**). Modeling these mutations onto the yeast 6PGDH structure, predicts both mutations to affect substrate and NADP^+^ binding as well as product release (16, 17) (**Fig. S1B-C**).

The *pgd3-umu1* null allele was discovered in color-converted W22 (15). The *pgd1-null* and *pgd2-125* alleles were crossed five times to color-converted W22 followed by three self-pollination generations to resynthesize the *pgd1; pgd2* double mutant. Loss of cytosolic 6PGDH activity was confirmed with isozyme activity assays of developing endosperm extracts (**Fig. 1A**). Although no PGD1/PGD2 activity was detectable, the double mutant kernels are normal (**Fig. 1B**). In comparison, *pgd3* mutants have severe grain-fill and defective embryo phenotypes in W22 (**Fig. 1C**).

**Figure 1.**
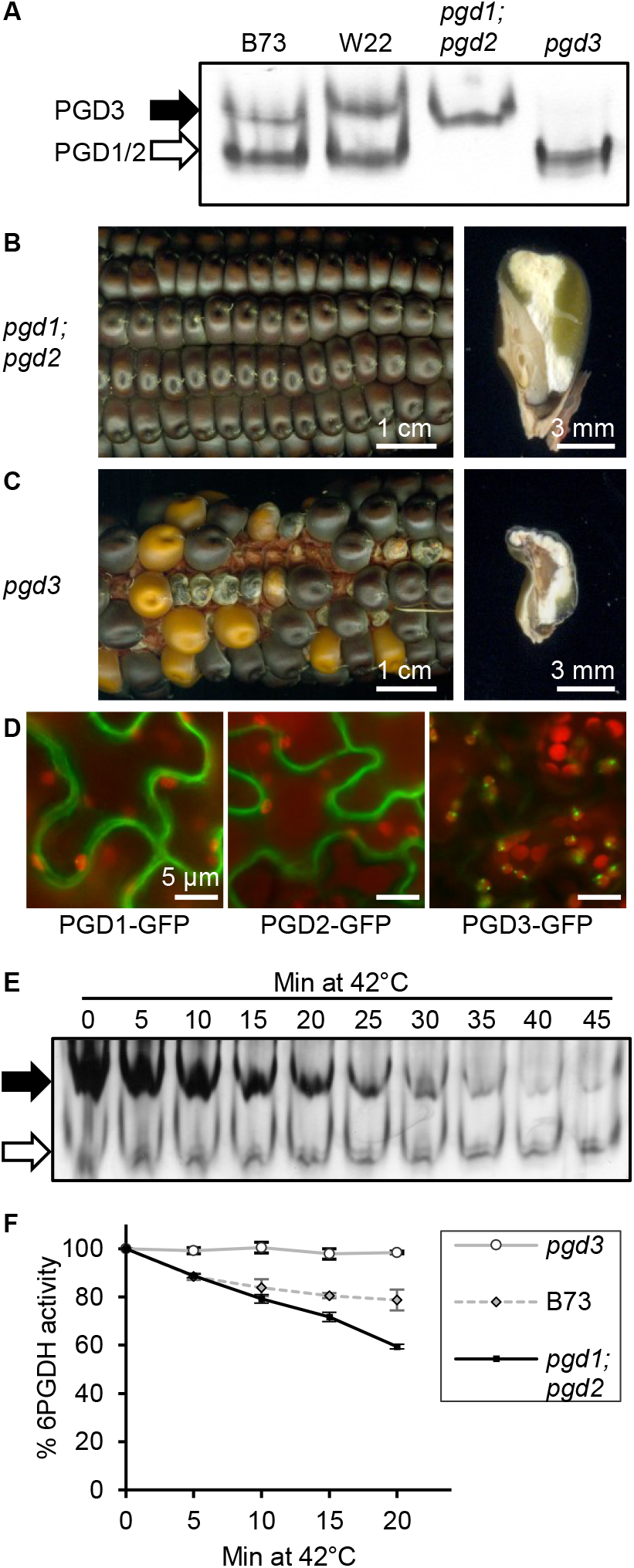
Plastid-localized 6PGDH is required for grain-fill and is heat sensitive. **(A)** Native PAGE of endosperm protein extracts stained for 6PGDH isozyme activity. B73 and W22 inbred lines show activity of all three isozymes with genetic variation for electrophoretic mobility. The black arrow indicates PGD3 homodimers. The white arrow indicates homodimers and the heterodimer of PGD1 and PGD2. **(B)** Homozygous *pgd1; pgd2* phenotypes in W22. Left panel is a mature homozygous double mutant ear. Right panel is a sagittal section. **(C)** Ear and sagittal kernel section phenotypes for *pgd3* in W22. Left panel is a *pgd3*/+ self-pollination. Right panel is a sectioned *pgd3* homozygous kernel. **(D)** *N. benthamiana* agroinfiltrated leaves expressing PGD1-GFP, PGD2-GFP, and PGD3-GFP fusion proteins. Merged images from spinning disk confocal fluorescence microscopy showing chlorophyll (red) and GFP (green) channels. **(E)** 6PGDH isozyme activity assay of endosperm extracts heated at 42°C. Extracts were transferred to ice until all treatments were completed. Black arrow indicates PGD3, and white arrow points to PGD1 and PGD2 activity. **(F)** Spectrophotometric assays for total 6PGDH activity of endosperm extracts heated at 42°C. B73 (diamonds, gray dashed) expresses all isozymes. *pgd1; pgd2* double mutants (squares, black solid) only have PGD3 activity, while *pgd3* mutants (circles, gray solid) only have PGD1 and PGD2 activity. Error bars indicate SD of three biological replicates.

Cytosolic localization of PGD1 and PGD2 has been inferred by cell fractionation, phylogenetics, and bioinformatics (11, 14, 17). To test whether PGD1 and PGD2 are exclusively cytoplasmic, PGD1 and PGD2 were fused in-frame with a C-terminal GFP for transient expression in *N. benthamiana* epidermis. Both PGD1 and PGD2 accumulate in the peripheral region of pavement cells, consistent with cytosolic localization (**Fig. 1D**). To the contrary, PGD3-GFP fusion co-localized with chloroplasts. Taken together these data confirm that PGD1 and PGD2 are localized to the cytosol and that loss of cytoplasmic activity does not impact the kernel visibly. Thus, plastid-localized 6PGDH is the sole 6PGDH isozyme required for maize kernel development.

### Plastidic 6PGDH activity is reduced by heat treatment

Long-term growth at elevated temperatures during grain-fill reduced the level of 6-phosphogluconate in endosperm (6) suggesting that flux through the PPP could be a heat sensitive step in endosperm metabolism. Heat stability of 6PGDH was tested by treating W22 endosperm extracts at 42°C and assaying isozyme activity by native PAGE (**Fig. 1E**). PGD3 activity progressively decreased as the length of the heat treatment increased. The PGD1/PGD2 activity band was unaffected. To quantify the decrease of PGD3 enzyme activity, endosperm extracts from B73, homozygous *pgd1; pgd2* double mutants in their reference genetic background, and homozygous *pgd3* in W22 were treated at 42°C. Total enzyme activity was measured spectrophotometrically (**Fig. 1F**). In *pgd3*, PGD1 and PGD2 isozyme activity remains, and this cytosolic activity is stable during heat treatment. Normal B73 extracts have a mix of PGD1, PGD2, and PGD3 isozymes and show a decrease of 25% activity during the heat treatment. The *pgd1; pgd2* mutant only has plastidic PGD3 activity, which decreases to 60% of starting activity after 20 min of heat treatment. Heat sensitivity of PGD3 is more apparent in embryo extracts, where total activity from the three isozymes in the normal embryos decreases by 60% after heat treatment (**Fig. S2**). In *pgd1; pgd2* mutant embryo extracts, PGD3 activity decreases by 80% during heat treatment. These results indicate that PGD3 activity is more sensitive to high temperatures, whereas PGD1 and PGD2 activity is unaffected by heat.

### Engineering heat stable amyloplastic 6PGDH isozymes

Heat stable 6PGDH was expressed in maize amyloplasts by targeting PGD1 and PGD2 to the organelle. Based on the yeast 6PGDH structure, the precise N- and C-termini of the peptides are likely required for correct folding and dimerization of the enzyme (16). Consequently, the N-terminal chloroplast targeting peptide of maize granule-bound starch synthase I (GBSS1), encoded at the *waxy1* (*wx1*) locus (18) was fused to the coding sequence of *Pgd1* and *Pgd2* by gene synthesis (**Fig. 2A**). Prior protein sequencing of mature GBSS1 identified the precise transit peptide processing site, which enables complete removal of the transit peptide after import into plastids (18, 19). The fused genes were named *Wpgd1* and *Wpgd2*.

**Figure 2.**
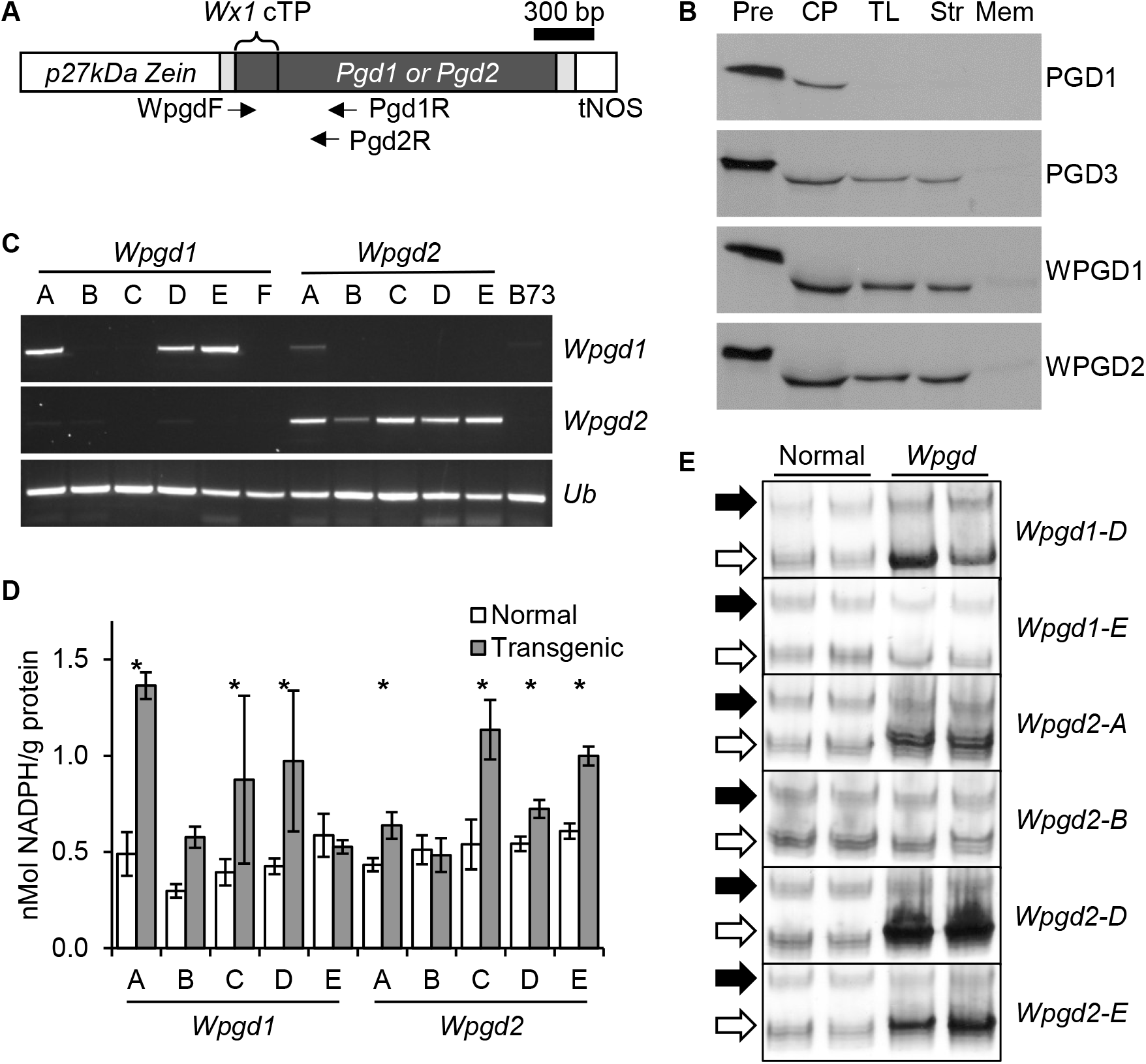
Recombinant *Wpgd* transgenes increase amyloplastic 6PGDH. **(A)** Schematic of recombinant *Wpgd* transgenes. The *Wx1* chloroplast transit peptide is fused to the full-length coding sequences of *Pgd1* or *Pgd2*. Promoter and terminator sequences are in white; UTRs are in grey; arrows indicate primers used for RT-PCR and genotyping. **(B)** *In vitro* chloroplast import of PGD1, PGD3, WPGD1, and WPGD2. Lanes are precursor (Pre) proteins, chloroplasts (CP) after the import assay, thermolysin (TL) treated chloroplasts after import, purified chloroplast stroma (Str) and membrane (Mem) after hypotonic lysis. **(C)** RT-PCR of endosperm RNA from transgenic kernels. Maize ubiquitin (*Ub*) was amplified as a loading control. **(D)** Endosperm 6PGDH enzyme activity for non-transgenic (normal) and transgenic sibling kernels. Error bars are SD and significant differences are indicated with an asterisk (p<0.05, Student’s t-test). **(E)** 6PGDH isozyme activity assays comparing individual normal and transgenic sibling endosperm from developing T_1_ kernels. Black arrows are PGD3 activity; white arrows are PGD1 and PGD2.

*In vitro* transcribed and translated proteins were tested for chloroplast targeting with import assays into purified pea chloroplasts (**Fig. 2B**). Wild-type PGD1 associated with chloroplasts but was outside of the outer envelope, as thermolysin digested the radiolabeled protein. By contrast, PGD3 was resistant to thermolysin treatment and fractionated as a soluble protein after hypotonic lysis and centrifugation of the chloroplast fractions. The WPGD1 and WPGD2 proteins followed the pattern of PGD3, indicating the engineered proteins import into plastids.

The *Wpgd1* and *Wpgd2* genes were cloned into the pIPK27-MCSBAR binary transformation vector with a 27 kDa γ-zein endosperm-specific promoter and transformed into the maize HiII line (**Fig. 2A**, **Fig. S3**). Expression of *Wpgd1* and *Wpgd2* was tested by RT-PCR in developing T_1_ seeds from self-pollinations of hemizygous T_0_ plants (**Fig. 2C**). In parallel, immature T_1_ kernels were characterized for 6PGDH activity. Embryos were genotyped for the transgene and endosperms were assayed for activity. Most transgenic events increased total 6PGDH enzyme activity (**Fig. 2D**), while native PAGE showed more activity in the faster migrating, PGD1/PGD2 band (**Fig. 2E**).

### *Wpgd* transgenes complement defective *pgd3* endosperm

If targeted correctly, WPGD1 and WPGD2 are expected to rescue the defective endosperm phenotype of *pgd3* mutants. Single hemizygous transgenics, either *Wpgd1/*- or *Wpgd2/-*, were crossed with *pgd3/+* plants. F_2_ kernels from *pgd3/+; Wpgd/−* are expected to be 75% normal due to segregation of a wild-type copy of the *pgd3* locus (**Fig. 3A**). The remaining 25% of *pgd3* mutants could show transgene rescue. Based on non-concordant kernels from B-A translocation uncovering crosses, *pgd3* mutants have independent endosperm and embryo phenotypes (14). Both *Wpgd1* and *Wpgd2* are expressed from an endosperm-specific promoter, which would not be expected to rescue the embryo lethal *pgd3* phenotype. Consequently, the transgenes are predicted to cause increased grain-fill without rescuing embryo development in 18.75% of F_2_ kernels from the total population, while *pgd3* mutant kernels should be reduced to 6.25% of total F_2_ kernels (**Fig. 3A**).

**Figure 3.**
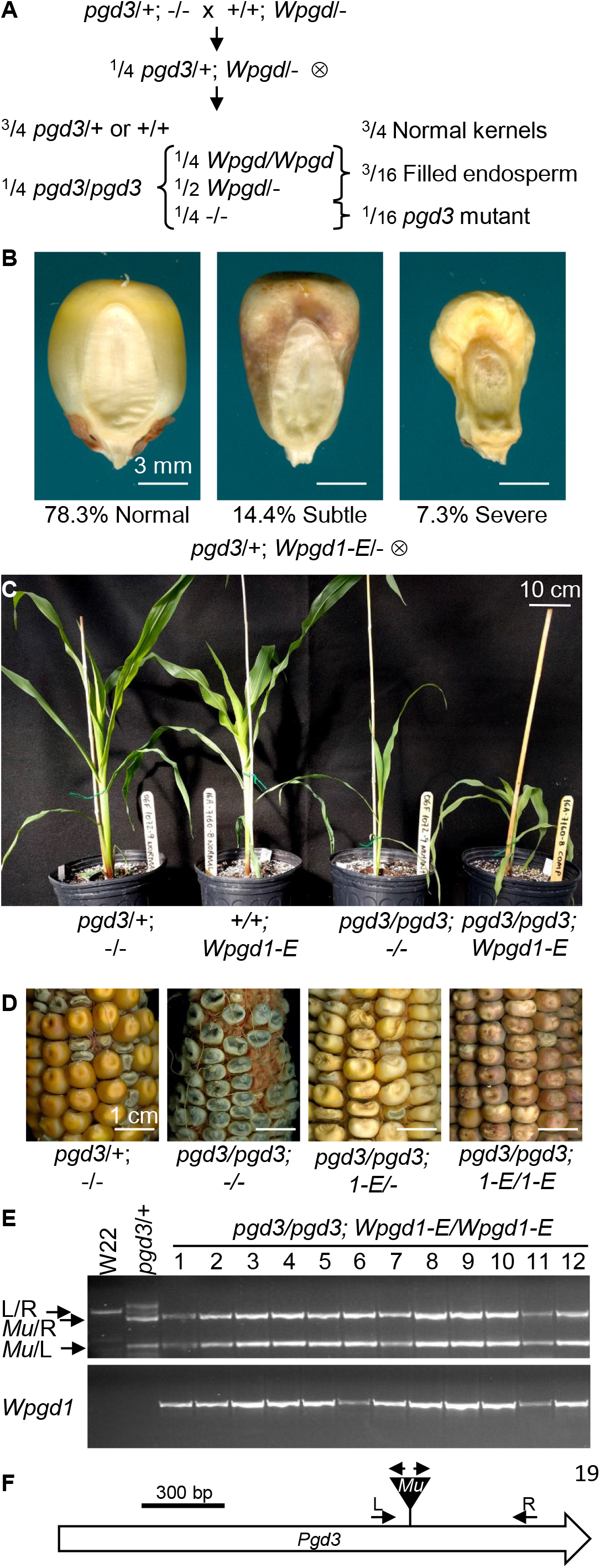
*Wpgd* transgenes rescue *pgd3* seed phenotype. **(A)** Schematic of crosses and expected phenotypes. Heterozygous *pgd3/+* plants were crossed with hemizygous *Wpgd/−* transgenics. F_1_ *pgd3/+; Wpgd/−* were self-pollinated and 75% of homozygous *pgd3* are expected to be rescued for endosperm grain-fill phenotypes. **(B)** Germinal view of representative kernels observed in F_1:2_ ears segregating for *pgd3* and the *Wpgd1-E* transgenic event. Frequencies are data from five ears. Scale bars are 3 mm. **(C)** Representative plants at 35 DAG comparing a *pgd3* heterozygote, a *Wpgd1-E* transgenic, a homozygous *pgd3* plant, and a homozygous *pgd3* with the *Wpgd1-E* transgene. **(D)** Self-pollinated ears comparing *pgd3* phenotypes in W22 and F_2:3_ ears from crosses with *Wpgd1-E*. Genotypes of the parent plants are given below each image. Transgenic genotypes were determined by PCR analysis of F_3_ kernels. Scale bars are 1 cm. **(E)** PCR genotyping of double homozygous *pgd3; Wpgd1-E* F_3_ kernels. **(F)** Schematic of the *Pgd3* locus and primers used for genotyping in panel E.

Consistent with this model of genetic function, we observed two classes of mutant kernels in *pgd3/+; Wpgd/−* self-pollinations (**Fig 3B**). Two transgenic events, *Wpgd1-F* and *Wpgd2-F* had severe *pgd3*-like mutants near a 6.25% frequency (**Table S1**). *Wpgd1-E* and *Wpgd2-G* had a higher frequency of severe mutants of 7.3% and 9.9%, respectively. Nevertheless, all four transgenes segregated for a subtle mutant phenotype with a large endosperm that frequently supported embryo development (**Table S1**; **Fig. 3B**, **Fig. S4A**). Combining the severe and subtle classes resulted in significantly fewer than 25% mutants with the subtle phenotype always having a frequency lower than 18.75% (**Table S1**). Contrary to the genetic model, the data suggest that the transgenes fully rescue a fraction of homozygous *pgd3* mutants to develop normal kernels.

Plant development was compared between embryo-rescued homozygous *pgd3* mutants and subtle F2 kernels from *pgd3/+; Wpgd/+* self-pollinations (**Fig 3C**, **Fig. S4B**). Normal siblings were controls for plant size and color. Consistent with endosperm-specific expression of the transgenes, both non-transgenic and transgenic *pgd3* mutants grow slowly and have a virescent, pale-green color in expanding leaves (**Fig 3C**, **Fig. S4B**). The mutant plants were fertile with non-transgenic *pgd3/pgd3* progeny developing all severe grain-fill phenotypes. Self-pollination of *pgd3/pgd3; Wpgd/−* plants develop F_3_ kernels with predominantly normal grain-fill and a subset of reduced grain-fill kernels lacking the transgene (**Fig. 3D**; **Fig. S4C**). By contrast, *pgd3/pgd3; Wpgd/Wpgd* double homozygotes develop nearly 100% normal F_3_ kernels (**Fig. 3D**; **Fig. S4C**). Genotyping F_3_ kernels from F_2_ transgenic *pgd3* mutants confirmed the inferred F_2_ plant genotypes. For example, all F_3_ kernels from normal ears were homozygous for *pgd3* and positive for the transgene (**Fig 3E**; **Fig S4D**). These results show that homozygous *Wpgd* transgenes can fully complement kernel development of *pgd3* mutants.

### *Wpgd* transgenes mitigate grain yield losses under heat stress

Three events for each of the *Wpgd1* and *Wpgd2* transgenes were selected for field trials under heat stress. Heat stability of 6PGDH activity was tested *in vitro* with *Wpgd1-C*, *Wpgd1-D*, *Wpgd2-C*, and *Wpgd2-D* showing increased activity relative to non-transgenic siblings (**Fig. 4A**). To test heat stress in the field, the six *Wpgd* transgenic events and non-transgenic segregants were planted on March 15, 2017 (planting 1) and April 12, 2017 (planting 2) in Citra, Florida. The field site is at a Florida Automatic Weather Network (FAWN) monitoring station that recorded data every 15 minutes throughout the growing seasons (**Fig. 4B**). Under controlled conditions, the W22 inbred shows reduced grain-fill under elevated nighttime temperatures (6). In these two plantings, the grain-fill period from first pollination until harvest showed no difference in the length of time at temperatures above 33°C (**Fig. S5A**). However, planting 2 experienced an increase in nighttime temperatures from 8-32 DAP with no time spent below 22°C, while planting 1 experienced an average of 4.5 h/night in cooler temperatures, <22°C, during this developmental window (**Fig. S5B**). Planting 2 had an overall reduction in yield parameters such as ear weight, grain yield per plant, and number of kernels per plant (**Fig. 4C-E**). Average ear weights of non-transgenics were 133 g in planting 1 and 80 g in planting 2. This is a 40% reduction of normal yield in planting 2, which is consistent with a high temperature threshold effect on maize yields (20, 21).

**Figure 4.**
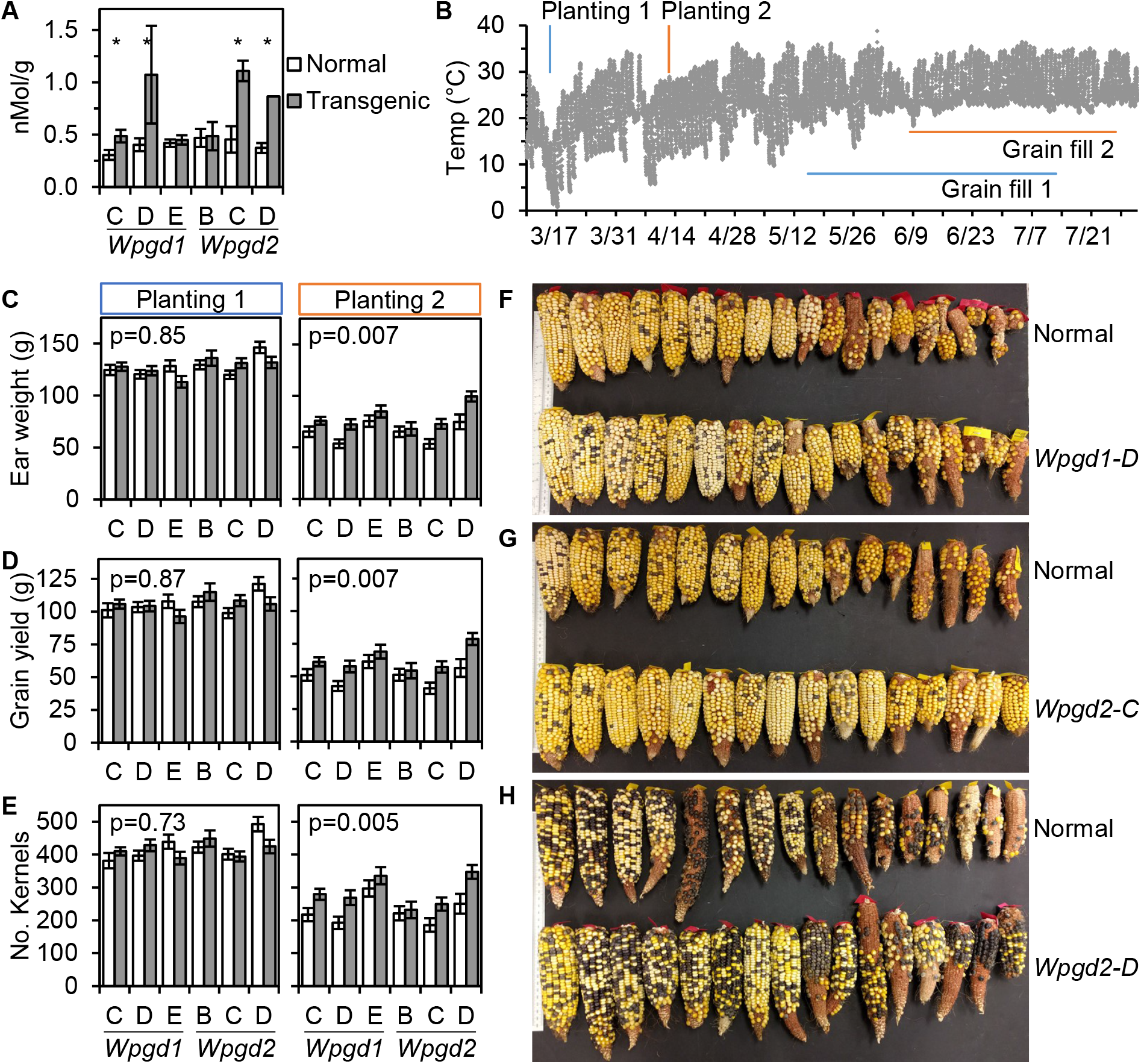
*Wpgd* transgenes mitigate grain yield losses due to heat stress. **(A)** Endosperm 6PGDH enzyme activity for normal (white) and transgenic (gray) sibling kernels after extracts were held at 50°C for 10 min. Error bars are SD. Asterisks indicate significant differences between transgenic and normal siblings (p<0.05 for Student’s t-test). **(B)** Air temperature at 60 cm for the 2017 Citra, FL field site. Readings are at 15 min intervals. Planting date and the pollination to harvest time period are indicated for both plantings. **(C-E)** Yield parameters of ear weight **(C)**, grain yield per plant **(D)**, and kernel number per plant **(E)** for normal (white) and transgenic (gray) field plots. Plot averages and standard errors are shown with the same y-axis scale for plantings 1 and 2. The p-value for paired t-tests of the normal and transgenic plots is given for each yield parameter. **(F-H)** Images of ears harvested from normal and transgenic plots in planting 2.

In planting 1, transgenic and non-transgenic plants showed no significant differences, while transgenics in planting 2 had significant increases in yield relative to non-transgenics. Yield increases were driven by an average increase of 25.6% in kernel number with corresponding increases in grain yield, ear weight, and cob weight, while ear length and 100 kernel weight were not significantly different (**Fig 4C-E**, **Fig S5C-E**). The overall increase in seed set can be seen as an increased number of ears with fuller kernel set and tip fill (**Fig 4F-H**). *Wpgd1-D*, *Wpgd2-C*, and *Wpgd2-D* all gave >35% increase in grain yield in the heat stress conditions. *Wpgd1-E* and *Wpgd2-B*, which had no significant increase in heat-stable 6PGDH activity *in vitro*, had the smallest effects on grain yield with a 12.4% and 6.1% increase in planting 2, respectively.

## Discussion

### Grain yield is impaired by heat-stress

High temperatures affect grain yield with up to 50% yield loss, which can be caused by a variety of mechanisms (22–27). Heat can reduce seed set by accelerating plant development, causing asynchrony between male and female gamete development as well as reducing ovary and pollen viability (28–30). Accelerated development during grain-fill reduces the time available for seed storage molecule accumulation (6). Heat also reduces gene expression of carbohydrate metabolic enzymes, potentially reducing the flux of fixed carbon into carbohydrate and protein storage (6, 31, 32). It is important to understand these mechanisms and how they interrelate in order to breed cereals that maintain yield in high temperature environments.

Our results support a direct heat-sensitivity of PGD3, while PGD1 and PGD2 are heat stable enzymes. The subcellular specialization of PGD3 activity provided the opportunity to test whether PGD3 heat sensitivity observed *in vitro* contributes to yield losses. Providing heat stable PGD1 or PGD2 to the endosperm amyloplast mitigated yield losses indicating that the heat sensitivity of PGD3 is a factor in the field. The primary yield improvement of *Wpgd1* or *Wpgd2* was due to increased kernel set rather than changes in grain-fill.

The first committed step of starch biosynthesis is catalyzed by ADP-glucose pyrophosphorylase (AGPase) and is a thermolabile enzyme (6). Increasing heat stability of AGPase in maize endosperm also results in mitigation of heat-induced yield losses (33, 34). Like thermostable 6PGDH, heat tolerant AGPases drive an increase in seed set rather than increasing seed weight. Kernel set could potentially be driven by early expression of the transgenes in developing seeds. However, enhanced AGPase genes appear to act through maternal tissues, because non-transgenic kernels fill at a higher rate on transgenic maternal plants (33, 34). Additional field trials will be needed to determine if the *Wpgd1* and *Wpgd2* transgenes act with a similar genetic mechanism.

### Subcellular specialization of carbon metabolism enzymes in cereals

Prior studies showed that 6PGDH isozymes in maize have distinct functions. PGD3 is critical for endosperm starch accumulation (14, 15). By contrast, the PGD1 and PGD2 isozymes do not affect seed phenotype (12, 13, 17). We further validated that PGD1 and PGD2 are cytosolic enzymes with GFP fusion proteins, and we showed that *pgd1; pgd2* double mutants have no seed phenotype in the W22 genetic background. Moreover, targeting PGD1 or PGD2 to the amyloplast was sufficient to rescue grain-fill in *pgd3* mutants. Therefore, the subcellular compartment localization of 6PGDH is the critical metabolic factor for kernel development. The complete rescue of *pgd3* mutants by endosperm-specific *Wpgd* transgenes suggest that this oxPPP step is most critical in endosperm.

In *Arabidopsis thaliana*, plastid-localized oxPPP has also been shown to be critical for endosperm and embryo development, although the relative importance of oxPPP in endosperm versus embryo was not investigated (35). Comprehensive analysis of Arabidopsis mutations removing individual steps of the plastid-localized PPP and related enzymes suggest that synthesis of ribose-5-phosphate and histidine are the primary requirements for seed development to proceed past the globular embryo stage (35). Metabolite profiling of maize *pgd3* mutant endosperm showed a moderate increase in histidine and less than two fold changes in pentose levels suggesting that these metabolite pools are much less impacted than would be predicted by Arabidopsis mutants (14). Intriguingly, metabolite profiling of heat-stressed maize endosperm shows that the substrate for 6PGDH, 6-phosphogluconate, is reduced while ribose products are increased suggesting a perturbation of oxPPP during heat stress that is inconsistent with specific loss of 6PGDH activity (6). However, this study also tested 20 carbon metabolism enzyme activities, and six were differentially affected by high nighttime heat supporting more complex heat stress impacts on carbon metabolism than simply reducing 6PGDH activity.

### Potential for translational improvements in breeding programs

Cereals evolved a predominant cytoplasmic localization of AGPase activity specifically in the endosperm (36–38).Transgenic modification of cytoplasmic AGPase improves seed set in both rice and wheat suggesting that the specialized carbon metabolism needed for endosperm grain-fill is a common metabolic feature to cereals (39–42). Throughout angiosperms, oxPPP enzymes are found in both cytoplasm and plastids, and the plastidic pathway is needed for seed development in both maize and Arabidopsis (10, 14). The plastid-localized 6PGDH represents a distinct phylogenetic group from cytoplasmic isozymes indicating that the genes encoding for plastid-targeted isozymes are most likely derived from a common ancestor, potentially the cyanobacterial endosymbiont (14). This evolutionary divergence raises the possibility that plastid-localized 6PGDH activity is heat-sensitive in many plants and could be a target for genetic improvement of heat tolerance.

This work illustrates the value of genetic engineering to generate and test novel traits that would be nearly impossible to identify by chance through traditional breeding methods. By defining a target trait of improved heat tolerance for plastid-localized 6PGDH, it is possible to develop germplasm with beneficial alleles in the oxPPP using conventional breeding or cisgenic insertions. For example, diverse germplasm could be screened for more heat tolerant alleles of *Pgd3*, which could then be incorporated into breeding programs. It is also possible to develop a cisgenic trait by inserting an exon encoding a chloroplast transit peptide in-frame with a cytosolic 6PGDH gene to develop a plastid-targeted heat-stable 6PGDH. These types of insertions have occurred over the course of evolution to give a mosaic chloroplast proteome of cyanobacterial and eukaryotic origin (43). Further field trials with hybrid *Wpgd1* and *Wpgd2* in diverse environments and germplasm are needed prior to significant efforts to incorporate novel heat-tolerant 6PGDH alleles in maize breeding germplasm.

## Materials and Methods

### Plant material

Maize was grown in the field at the University of Florida Plant Science Research and Education Unit in Citra, FL, or in a greenhouse at the Horticultural Sciences Department in Gainesville, FL. The *pgd3-umu1* allele and *pgd1-null; pgd2-125* double mutant were previously reported (13, 14). The *pgd1-null; pgd2-125* double mutant was crossed to the color-converted W22 inbred five times with double heterozygotes selected using PCR markers. Three self-pollination generations were used to select *pgd1-null; pgd2-125* double mutants in W22. The *Wpgd1* and *Wpgd2* constructs were transformed into the HiII genotype using *Agrobacterium tumefaciens* at the Iowa State University Plant Transformation Facility (http://www.agron.iastate.edu/ptf/). Transgenic plantlets were grown in a greenhouse at the University of Florida for crosses to B73 or *pgd3/+* plants.

### Molecular markers for *pgd1* and *pgd2*

The *pgd1-null* and *pgd2-125* alleles were sequenced from the homozygous double mutant stock genomic DNA. Products spanning the open reading frame of each gene were amplified with a high-fidelity Phusion® (NEB) DNA polymerase. PCR product sequences were compared with full-length cDNA sequences from W22 and B73 to identify mutations using the software ProSeq3.5 (44). Co-dominant markers were designed for genotyping the *pgd1-null* and *pgd2-125* mutations. The *pgd1-null* allele could be detected with an insertion-deletion (InDel) product using the primers: PGD1C-F: CTACGAGAGGGTCGACATGC and PGD1C-R: TGCAGGAAATCTCATTACCG. The *pgd2-125* allele was detected with a cleaved amplified polymorphic sequence (CAPS) marker using the primers: PGD2C-F: GCATCAAGAAGGCGTACGAT, PGD2C-R: TTACTCGACACGGTGGCATA. The product was digested by BsmFI (NEB) to detect mutant and normal alleles. Maize predicted proteins were modeled to the crystal structure of *Saccharomyces cerevisiae* Gnd1 (PDB:2P4Q) with Phyre2 (45).

### Subcellular Localization

RNA was extracted from B73 and W22 leaves using Trizol (ThermoFisher) and DNaseI (NEB) following the manufacturers’ protocols. Full-length cDNA for PGD1 and PGD2 were synthesized with the SuperScript® III First-Strand Synthesis System (ThermoFisher). *Pgd1* and *Pgd2* cDNA was amplified and cloned into a TOPO Zero Blunt® (ThermoFisher) vector (46, 47). The cDNA sequences were compared with the B73_v3 reference genome at MaizeGDB (48) and Gramene (49) using the software ProSeq3.5 (44). Open reading frames (ORFs) of *Pgd1*, *Pgd2*, and *Pgd3* were subcloned into the pENTR vector (Gateway ThermoFisher) (50), and then recombined into the binary vector pB7-MP:GFP (51). The PGD1-GFP, PGD2-GFP, and PGD3-GFP vectors were transformed into *Agrobacterium tumefaciens*. *Nicotiana benthamiana* was grown in a growth chamber at 22-24°C with 16/8 h day/night. Agrobacterium (OD_600_ = 0.6) was infiltrated into leaves of 4-week-old plants using a needleless syringe as described by (52). Fluorescence in epidermal cells of *N. benthamiana* leaves was visualized by spinning disk confocal microscopy (X81-DSU-Olympus) 48 h after transient transformation.

### Isozyme activity

Frozen tissues from total or dissected kernels were ground in liquid nitrogen and cold extraction buffer (100 mM Tris-HCl pH 7.5, 30 mM 1,4-Dithiothreitol (DTT), 15% (v/v) glycerol). Protein samples were separated by 10% native polyacrylamide gel electrophoresis (PAGE) at 30 mA for 2.5 h at 4°C. 6PGDH activity was assayed by incubating gels at room temperature for 30 min in the dark with 6PGDH staining solution (0.1 mg/mL NADP^+^, 0.1 mg/mL nitro blue tetrazolium, 0.1 mg/mL phenazine methosulfate, 0.5 mg/mL 6-phosphogluconate, 100 mM Tris-HCl, pH 7.5, and imaging as described (12).

### Total Enzyme Activity

Spectrophotometric determination of total 6PGDH was adapted from (53). Total protein was extracted with 300 μL of cold extraction buffer that contained, 50 mM HEPES (pH 7.5), 200 mM KCl, 10 mM MgCl_2_, 2.5 mM EDTA (pH 7.5) and 5% sucrose. Extracts were centrifuged at 1,600 g for 20 min at 4°C and desalted with Zebacolumns (ThermoFisher). Protein concentration was measured using a commercial Bradford assay (Bio-Rad). 6PGDH activity was detected with 0.1 mM NADP^+^, 0.1 mM 6-phosphogluconate, 0.2 mM Tris-HCl, 0.5 mM MgCl_2_. Spectrophotometric absorbance at 340 nm was measured every minute for 10 min. Absorbance was regressed against time, and the slope was used to determine enzyme activity in Units/mg protein from the Lambert-Beer Law.

### *In vitro* protein heat stability

Protein extracts were aliquoted into 10 or 20 μL volumes and placed in a 42°C or 50°C water bath. Control aliquots were kept in ice during the heat treatment. Heat-treated aliquots were placed in ice at 5 min intervals. 6PGDH enzyme activity was assayed from control and heat-treated protein extracts at the end of all heat treatments.

### *In vitro* chloroplast import

The *Wx1* transit peptide translational fusions with *Pgd1* and *Pgd2* were synthesized by GenScript. The *Wpgd1* and *Wpgd2* fusion genes as well as *Pgd1* and *Pgd3* ORFs were cloned in pGEM3Z (Promega). Chloroplast import assays were performed essentially as described by (54). Briefly, intact pea chloroplasts were isolated from *Pisum sativum* seedlings. *Pgd1, Pgd3, Wpgd1*, and *Wpgd2* were *in vitro* transcribed using SP6 polymerase (Promega) and translated using wheat germ extracts in the presence of [^3^H]-leucine. Diluted translation products were incubated with intact chloroplasts (0.33 mg chlorophyll/mL) and 5 mM Mg-ATP (Sigma) for 20 min in a 25°C water bath under 120 μE/m^2^/s light. After import, intact chloroplasts were purified, re-suspended, and treated with thermolysin (Sigma). Protease digestion was stopped with EDTA, and chloroplasts were purified. Chloroplast pellets were lysed by re-suspending in a hypotonic solution for 10 min, followed by the addition of one volume of 2x import buffer. Plastid lysates were centrifuged at 150,000 g for 20 min at 4°C to separate soluble and membrane proteins. Protein samples from the assay were separated with 10% SDS-PAGE followed by fluorography.

### Transgene constructs and molecular markers

*Wpgd1* and *Wpgd2* ORFs were sub-cloned in the binary vector pIPK-27-MCSBAR (**Fig. S3**), adapted from pIPK006 (55). The Gateway system cassette was replaced with a custom multiple cloning site, and the coding region and 35S terminator of the hygromycin resistance gene in the original plasmid was replaced by the bialophos resistance gene and the transcription termination region of the soybean *vspB* gene, respectively. The promoter of the maize 27 kDa zein gene promoter was introduced upstream of the multiple cloning site, comprising the genomic region from 1041 bp upstream to 60 bp downstream of the transcription start site of *zp27*, gene model GRMZM2G138727. The presence of the transgene was determined by resistance to glufosinate-ammonium and PCR of each construct specifically with the following primers: ZWPGD1F, AAATAGGCCGGAACAGGAC, PGD1R, ACAGAGATGGGGAACCCTTT, ZWPGD2F, AAACTGAGCCACGCAGAAGT, PGD2R, CTTGGAGGTCGTCCTGTTGT.

### Transgene rescue of *pgd3*

Heterozygous *pgd3-umu1/+* plants were crossed with *Wpgd2/-* and *Wpgd1/-* HiII T0 plants. F1 progeny from the crosses were evaluated for glufosinate resistance, self-pollinated, and genotyped for the transgenes and *pgd3*. The *pgd3* locus was genotyped with a codominant marker using the PGD3L, PGD3R, and Tir5 primers as described (15). F_2_ kernels from each ear were separated according to visual phenotype. Segregation ratios were evaluated with χ^2^ tests for goodness of fit using the models in **Table S1**. F_2_ plants were grown in the greenhouse, genotyped for *pgd3*, the presence of the *Wpgd* transgenes, and self-pollinated. F_3_ kernels were evaluated visually and were genotyped for segregation of *pgd3* and *Wpgd* transgenes by PCR. Kernel and ear images were acquired with a flatbed scanner, and plant images were taken with a digital camera.

### Field trial

For *Wpgd1-C*, *Wpgd1-D*, *Wpgd2-B*, and *Wpgd2-C*, BC_1_S_1_ kernels from B73 introgressions were selected from transgenic and non-transgenic sibling BC_1_ plants. BC_1_S_1_ kernels from a W22 introgression was selected for *Wpgd2-D*, while F_3_ kernels from a *Wpgd1-E* cross with W22 were selected from transgenic and non-transgenic sibling F_2_ plants. Replicated nurseries were planted on March 15, 2017 and April 12, 2017 at the Plant Science Research and Education Unit in Citra, Florida. Each nursery was separated into transgenic and non-transgenic blocks separated by a male sterile planting to reduce transgenic pollen contamination in non-transgenic controls. Each block had three randomized sub-blocks with a replicate of each genotype. Individual 6 m plots were planted with 30 kernels. In the transgenic blocks, plants were painted with a 2% glufosinate-ammonium (Bio-world), and non-transgenic segregants were removed (56). All plots were thinned to 20 plants per plot and allowed to open pollinate. Weather data were downloaded for the Citra field site at the FAWN database (https://fawn.ifas.ufl.edu/). Ears were harvested at maturity, dried at 40°C for 1 week, and stored at 10°C in 50% relative humidity until measured for ear length, ear weight, grain weight, cob weight, kernel number, and 100 kernel weight. Traits were averaged for sub-block replicates of the same genotype. Average trait values for transgenic and corresponding non-transgenic plots were analyzed by paired Student’s t-tests assuming a two-tailed distribution.

## Acknowledgments

We thank L. Curtis Hannah, Michael McCaffery, Yubing Li, Chi-Wah Tseung, and John Baier for discussions and technical assistance. This work was supported by National Institute of Food and Agriculture projects: 2011-67003-30215, 2018-51181-28419, and FLA-HOS-005468; Brazilian Conselho Nacional de Desenvolvimento Cientifico e Tecnológico pre-doctoral fellowship award 209426/2013-6; and the Vasil-Monsanto Endowment.

---

**Figure S1.**
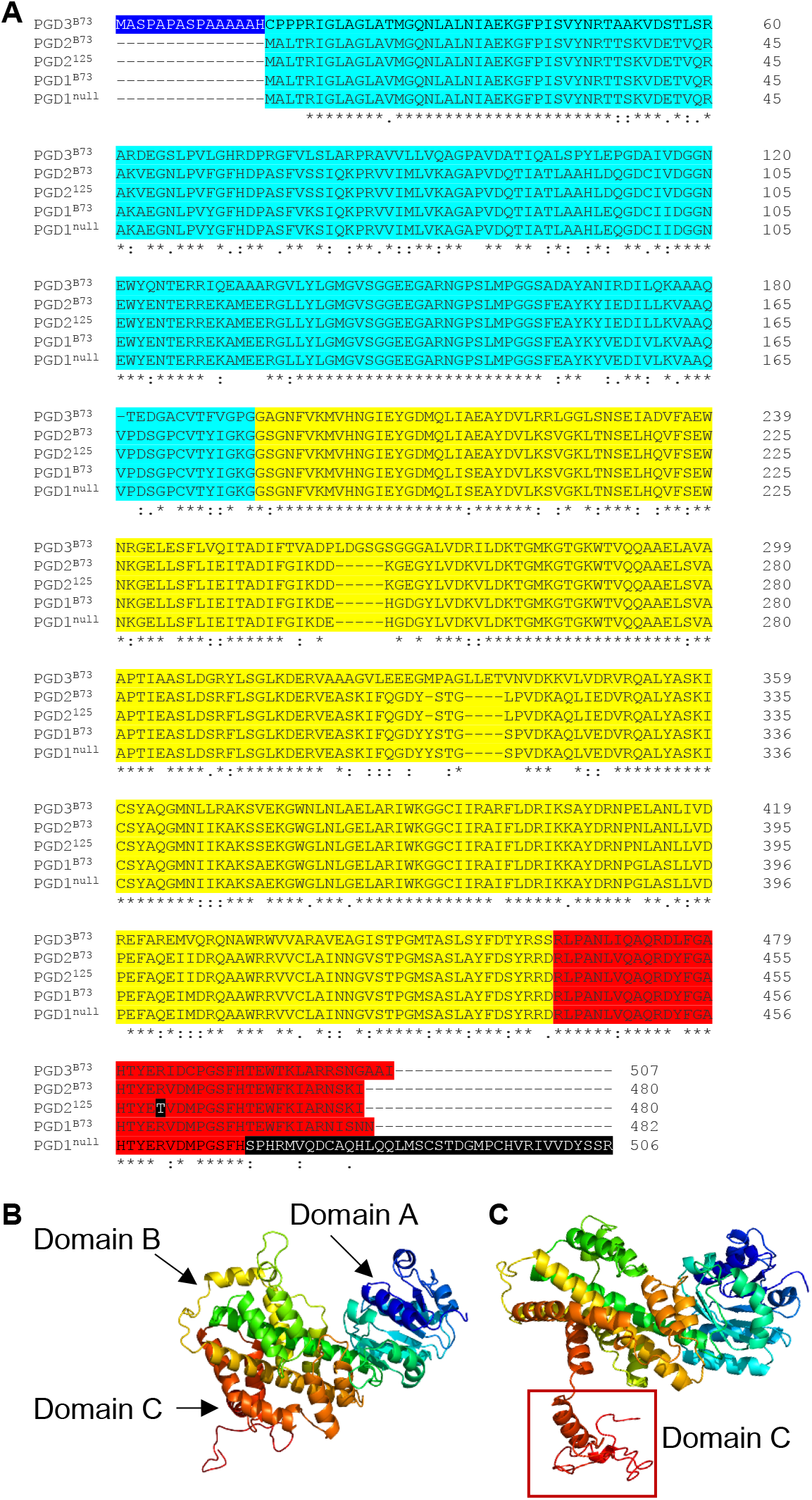
Predicted protein sequences for *pgd1-null* and *pgd2-125*. **(A)** Protein alignment of B73 6PGDH isozymes with predicted proteins from *pgd1-null* (PGD1^null^) and *pgd2-125* (PGD2^125^). Dark blue highlights the N-terminal putative transit peptide from plastidic 6PGDH (PGD3). Light blue highlights Domain A, the N-terminal coenzyme binding domain. Yellow highlights Domain B, the substrate binding and dimerization domain. Red highlights Domain C, the C-terminal tail that forms a lid on the substrate binding pocket. The 37 aa PGD1^null^ insertion and the PGD2^125^ R460T mutations are highlighted in black. **(B-C)** Predicted protein structures for PGD1^B73^ **(B)** and PGD1^null^ **(C)** modeled onto the crystal structure of *Saccharomyces cerevisiae* Gnd1 with Phyre2. The altered Domain C is indicated by a rectangle.

**Figure S2.**
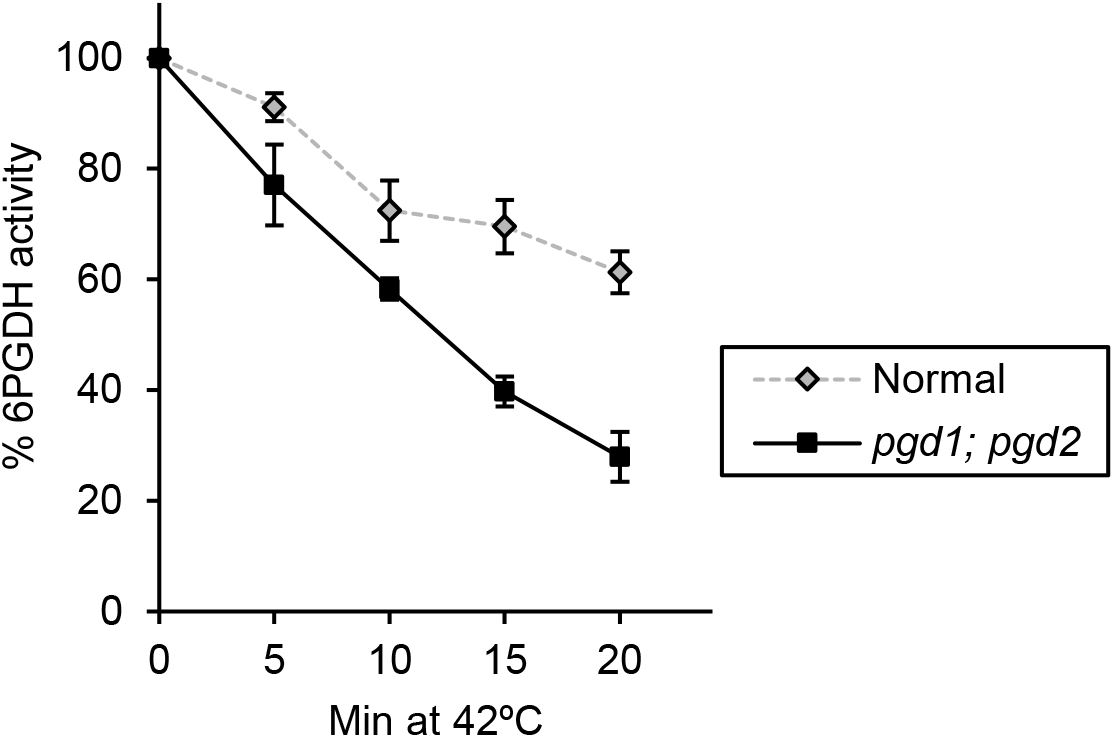
Spectrophotometric assays for total 6PGDH activity of embryo extracts heated at 42°C. Normal (diamonds, gray dashed) expresses all isozymes, while *pgd1; pgd2* double mutants (squares, black solid) only have PGD3 activity. Error bars indicate SD of three biological replicates.

**Figure S3.**
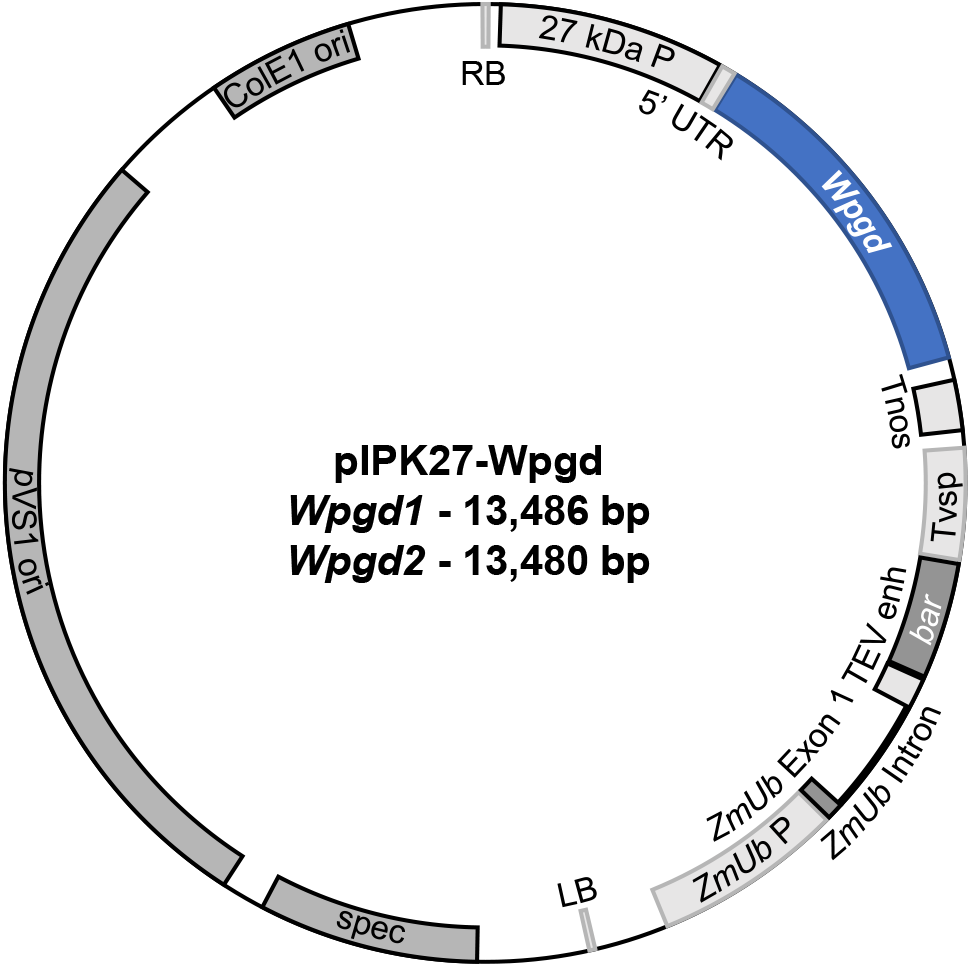
Schematic of *Wpgd* binary constructs used for plant transformation. Transformed DNA is between the right border (RB) and left border (LB). The 27 kDa zein promoter provides endosperm specific expression of the *Wpgd* fusion ORF. Transformed components of the vector are the nopaline synthase terminator (tNOS), the Tvsp terminator from *Glycine max*, the *Streptomyces hygroscopicus bar* gene encoding herbicide resistance to Basta/bialophos/phosphinothricin, the tobacco etch potyvirus enhancer (TEV enh), the *Zea mays ubiquitin* (*ZmUb*) promoter, exon 1 and intron 1.

**Supplemental Table 1.** F_2_ kernel phenotypes from *Wpgd* transgene rescue crosses with *pgd3* heterozygotes

| Transgenic event | Kernel phenotypes | | | $\chi^2$ p-values for segregation models | | |
| --- | --- | --- | --- | --- | --- | --- |
|  | Normal | Subtle | Severe | 3:1 | 4:3:1 | 15:1 |
| no transgene | 930 | - | 299 | 0.59 | N/A | N/A |
| *Wpgd1-E* | 1980 | 363 | 184 | 9.9x10<sup>-5</sup> | 4.1x10<sup>-8</sup> | 0.032 |
| *Wpgd1-F* | 2973 | 641 | 222 | 3.4x10<sup>-4</sup> | 1.5x10<sup>-3</sup> | 0.24 |
| *Wpgd2-F* | 506 | 74 | 45 | 5.8x10<sup>-4</sup> | 5.1x10<sup>-5</sup> | 0.33 |
| *Wpgd2-G* | 913 | 112 | 104 | 5.3x10<sup>-6</sup> | 1.7x10<sup>-15</sup> | 3.9x10<sup>-5</sup> |

<sup>a</sup>3:1 model tests subtle plus severe classes represent all *pgd3/pgd3* progeny
<sup>b</sup>4:3:1 model tests subtle are *pgd3*; *Wpgd* and severe are *pgd3* without a transgene.
<sup>c</sup>15:1 model tests if the frequency of severe fits expected *pgd3* without a transgene.

**Figure S4.**
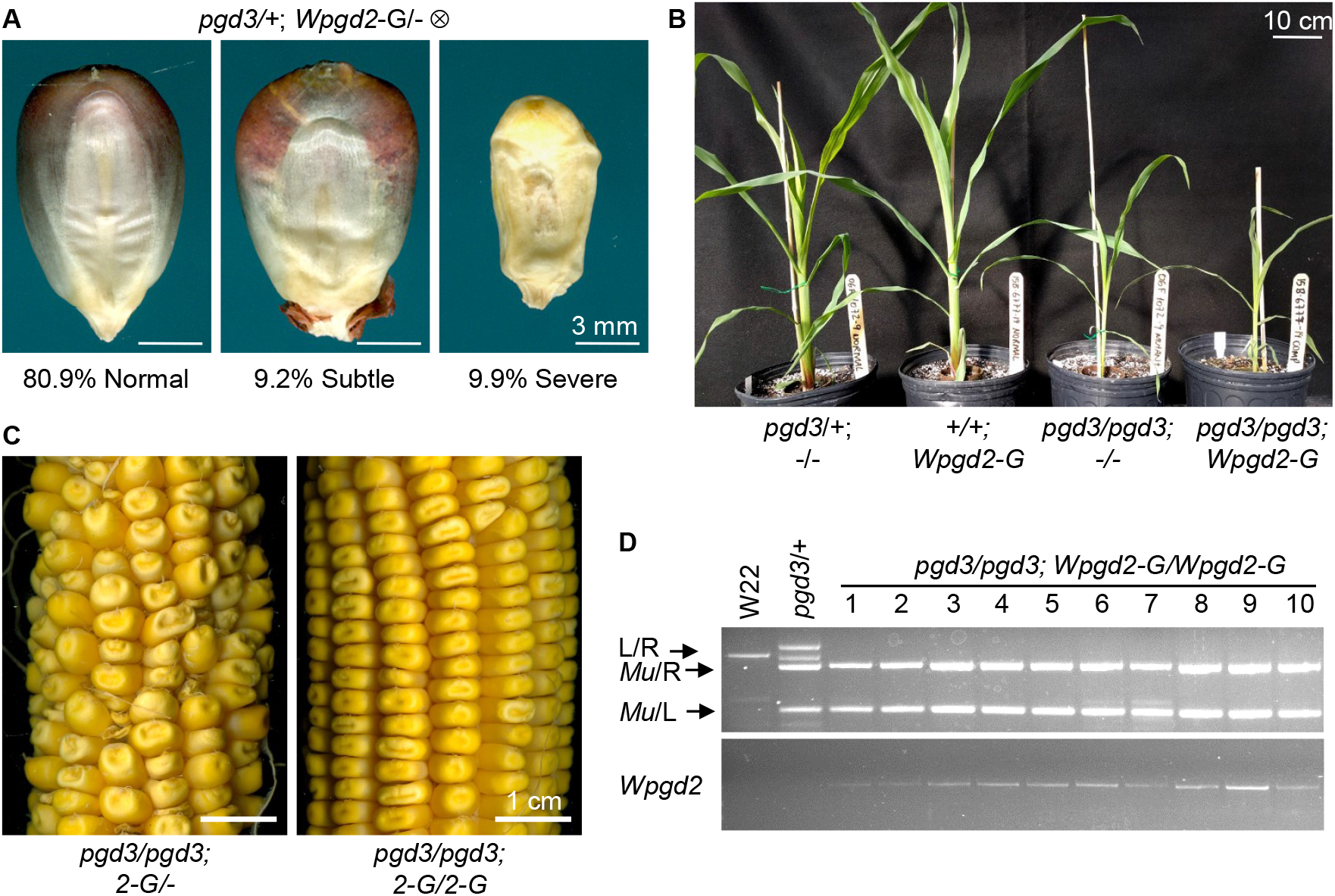
**(A)** Germinal view of representative kernels observed in F_2_ populations segregating for *pgd3* and the *Wpgd2-G* transgenic event. Frequencies are data from three F_1:2_ ears. Scale bars are 3 mm. **(B)** Representative plants at 35 DAG comparing a *pgd3* heterozygote, a *Wpgd2-G* transgenic, a homozygous *pgd3* plant, and a homozygous *pgd3* with the *Wpgd2-G* transgene. **(C)** Self-pollinated F_2:3_ ears. Genotypes of the F_2_ parent plants are given below each image. Transgenic genotypes were determined by PCR analysis of F_3_ kernels. Scale bars are 1 cm. **(D)** PCR genotyping of double homozygous *pgd3; Wpgd2-G* F_3_ kernels. PCR primers are the same as in Figures 1 and 3.

**Figure S5.**
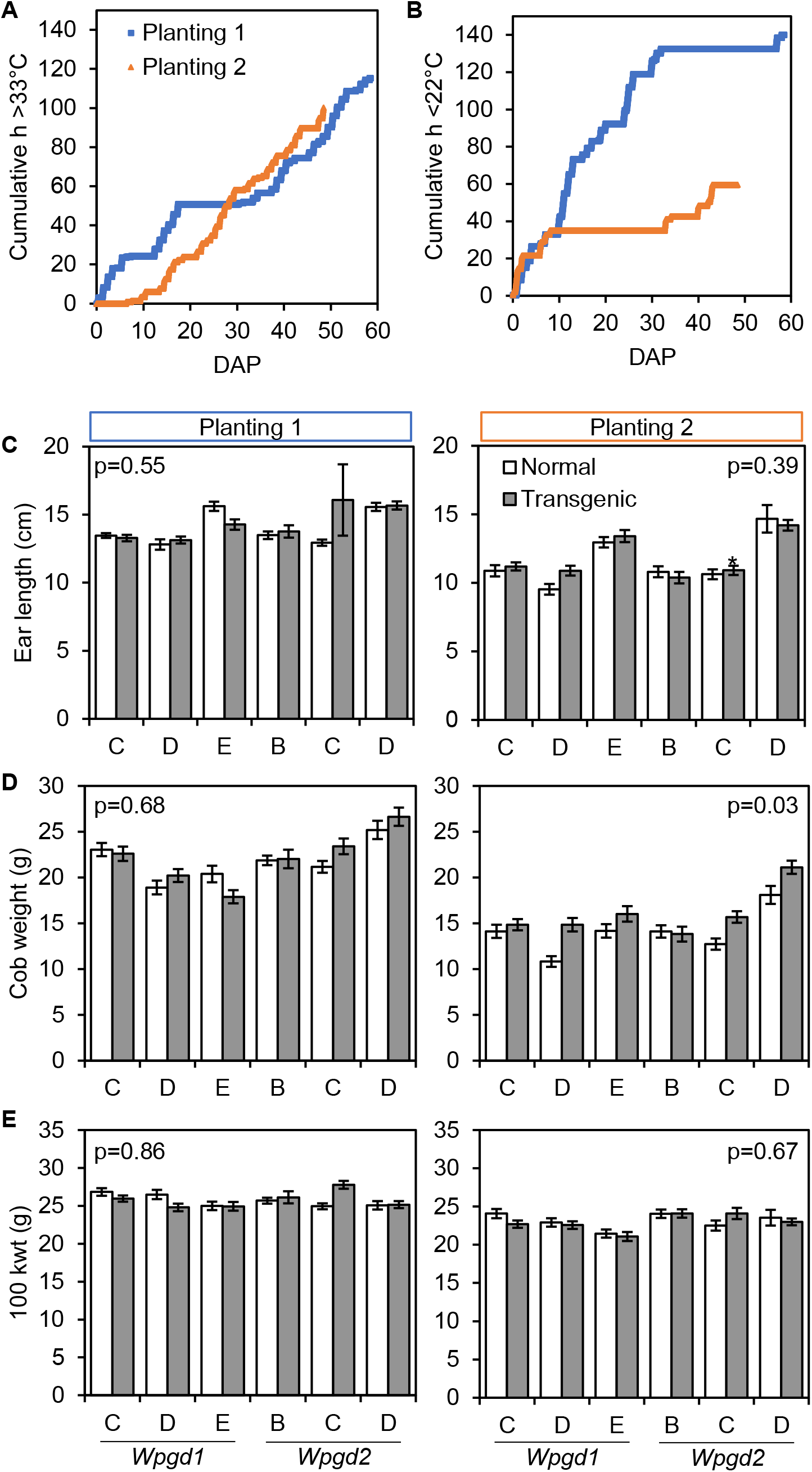
Additional weather data and yield traits from the 2017 field trial. **(A-B)** Cumulative time **(A)** above 33°C and **(B)** below 22°C during kernel set and grain-fill. X-axis shows days after pollination (DAP). Planting 1 is in blue and planting 2 is in orange. **(C-E)** Yield traits for **(C)** ear length, **(D)** cob weight, and **(E)**100 kernel weight, kwt. Each bar chart has plot averages and standard errors for normal (white) and transgenic (gray) field plots. Graphs for planting 1 and planting 2 have the same y-axis scale, and p-values are reported for paired t-tests of the average values for each normal-transgenic pair.

## Notes

The authors declare that they have no competing financial interests.

### Competing Interest Statement

The authors have declared no competing interest.

